# Animacy modulates gender agreement computation in Hindi: An ERP study

**DOI:** 10.1101/764811

**Authors:** Shikha Bhattamishra, R. Muralikrishnan, Kamal Kumar Choudhary

## Abstract

Gender has been investigated in psycholinguistics research both with respect to how it is retrieved when a noun is encountered and the computation of its agreement dependencies between different elements in a sentence. The literature is abound with differences as to the processing of gender agreement computations across languages and constructs. Similarly, neurophysiological evidence with respect to gender retrieval mechanisms differ wherein languages differ in the way the gender feature is retrieved when a noun is encountered based on different properties. The present study aims to scrutinize gender agreement computations in verb-agreement relations in Hindi and to ascertain if the intrinsic property (i.e. the animacy) of a noun participating in the relation plays a role in processing agreement dependencies. In addition, it also looks into the issue of whether the gender retrieval mechanism in the language influences agreement comprehension. Results revealed a positivity for the animate subject agreement and a negativity for the inanimate subject agreement violations at the position of the verb respectively. The results are suggestive of distinct processing mechanisms underlying semantic and syntactic gender. Further, the results also suggest that gender agreement computation in Hindi is modulated by the way gender is retrieved at the position of the noun.

## 1. Introduction

Neurophysiological studies on the processing of agreement constitute an essential part of the experimental work in the domain of sentence processing. Along with person and number, gender makes up the triad of the different features along which agreement dependencies are computed. Gender agreement relations may exist between a variety of constituents (adjective-noun, determiner-noun, subject/object-verb to speak a few) of a sentence in various languages of the world. Event-Related brain Potentials (ERPs) have been widely employed to study gender agreement in online language comprehension, and have focused, amongst other things, on investigating whether computing and comprehending gender agreement is a part of semantic or alternatively syntactic processing (Hagoort and Brown, 1999; Osterhout and Mobley, 1995); on uncovering the electrophysiological correlates of gender agreement as compared to that of agreement with other features i.e. number or person (Nevins, Dillon, Malhotra and Philips, 2007; Popov and Bastiaanse, 2018); on examining how gender agreement is established between certain constituents in a particular language (Barber and Carrerias, 2005); and on whether the electrophysiological signatures of gender agreement vary amongst different constituents (Barber and Carrerias, 2005; Gunter, Friederici and Schrifers, 2000). Except a handful of studies that examined verb-argument gender agreement dependencies (Deutsch and Bentin, 2001; Nevins et al.2007; Choudhary, 2011; Bornkessel-Schlesewsky, Choudhary, Witzlack-Makarevich and Bickel, 2008), most others have investigated adjective-noun and determiner-noun gender agreement (Hagoort, 2003; Aleman Banon, 2010; Martin Loeches, Nigbur, Casado, Hohfeld and Sommer, 2006; Popov, 2017). Besides these major strands of research on gender agreement, there have been few other issues in this domain that are slowly gaining prominence. One such issue concerns unearthing the electrophysiological correlates of the two different types of gender, namely syntactic gender and semantic gender^1^. Studies in this sub-domain of gender agreement (mostly investigating modifier-head relations) have not been conclusive on whether the underlying processing mechanism is sensitive to such a distinction. A point to be noted here is that the critical position in most of these studies is the noun. This also brings up another line of research on gender that is closely related to gender agreement. This field of research focuses on the retrieval mechanisms of the gender of a noun. Primarily, there seem to be two schools of thought: one that postulates a single route for the gender retrieval of a noun (Jescheniak and Levelt, 1994; Roelofs, 1992) and the other, which proposes a dual-route mechanism^2^(Gollan and Frost, 2001). Results in this regard are inconclusive, with certain studies finding evidence for the presence of a dual-route mechanism, while others do not. What is more intriguing is that these studies do not find evidence for the gender retrieval mechanism influencing agreement computation. However, if one looks at it closely, one would find that again the critical point in these studies is the noun. Based on the literature available in these major disciplines of experimental research on gender agreement and gender retrieval mechanism, it becomes clear that more work is needed to probe the gender agreement relations between a verb and its argument. This is with respect to both the less explored syntactic-semantic gender divide and the gender retrieval mechanism to check if it influences the computation of agreement dependencies between these constituents.

The present study sets out to fill this lacuna by investigating gender agreement relations between a verb and its subject argument in an intransitive sentence in Hindi, an SOV language spoken mainly in the Indian subcontinent. The choice of the language is imperative since it allows for the finite verb to agree with the highest nominative argument in a sentence (Kachru, 2006). Further, both animate as well as inanimate nouns can be the subject in an intransitive construction in Hindi. Thus the syntactic-semantic gender divide can be investigated in the experiment by manipulating the animacy of the subject noun. Also, by making the verb the critical position in the experiment, the study sets out to scrutinize the question of whether or not the different ways of retrieving the gender of a noun influence gender agreement dependency computation. Since both in the syntactic-semantic divide and in the issue of different gender retrieval mechanisms, animacy forms the basis of the division, the overarching goal of this experiment was to ascertain if the intrinsic properties of a noun such as animacy impacts the comprehension of agreement in a sentence. Though there have been previous works in the language (for instance, Nevins et al., 2007; Choudhary, 2011; Bornkessel-Schlesewsky et al., 2008), the line of research in those studies was quite different. While Nevins et al. (2007) probed into the ERP correlates of gender, number and person agreement between a subject and a verb, Choudhary (2011) and Bornkessel-Schlesewsky et al. (2008) investigated the electrophysiological correlates of long distance agreement (LDA) in the language. Thus, the current work has multiple goals: (i) to identify the ERP correlates of gender agreement between a finite verb and its subject argument in an intransitive sentence (ii) to examine if semantic and syntactic gender behave differently or in a similar manner (iv) to ascertain if the gender retrieval mechanism of a noun has any ramifications on the gender agreement computation mechanism and (iv) since animacy forms the basis of many of these divisions discussed above, to bring to light if the animacy of a noun influences the way gender agreement is computed between a verb and its subject noun.

## 2. Neurophysiological evidences on gender agreement and retrieval

### 2.1. Neurophysiology of Gender Agreement

Gender agreement from a comprehension point of view has been looked into in varied languages across the world (*Dutch:* Hagoort and Brown, 1999; Hagoort, 2003; Popov and Bastiaanse, 2018; *Spanish:* Barber, Salillas and Carrerias, 2004; Wicha, Moreno and Kutas, 2004; Barber and Carrerias, 2005; Martin-Loeches et al., 2006; Guajardo and Wicha, 2014; Aleman-Banon, 2016; *Italian:* Molinaro, Vespignani and Job, 2008; Popov, 2017; *English:* Osterhout and Mobley, 1995; Osterhout, 1997; *Hebrew:* Deutsch and Bentin, 2001; *German:* Gunter et al., 2000; *Hindi:* Nevins et al., 2007; Choudhary, 2011; Bornkessel-Schlesewsky et al., 2008). A LAN-P600 pattern (Hinojosa, Martin-Loeches, Casado, Munoz and Rubia, 2003; Barber and Carrerias, 2005; Gunter et al., 2000) or a lone P600 effect (Osterhout and Mobley, 1995; Popov and Bastiaanse, 2018) has been reported in the literature with respect to most studies investigating agreement violations. The N400, though relatively less common than the LAN, is also a part of the repertoire of electrophysiological studies on gender agreement (Wicha et al., 2004; Guajardo and Wicha, 2014; Popov, 2017, Deutsch and Bentin, 2001). The LAN has mostly been attributed to detection of a morphosyntactic mismatch^3^. The P600 found in studies of agreement processing has been mostly attributed to repair and reanalysis processes (Friederici, 2002; Osterhout and Holcomb, 1992) or well-formedness issues (Bornkessel and Schlesewsky, 2006). The N400 has been mainly attributed to the processing of conceptual information with respect to semantic gender.

As pointed out earlier, one of the lines of research on gender agreement concerns scrutinizing if the two facets of gender, i.e. semantic or syntactic gender, are treated similarly by the processing mechanism. One of the initial studies in this respect was done by Deutsch and Bentin (2001) who examined the agreement relation between a predicate and its subject argument manipulating the animacy of the subject argument (e.g. The woman saw that the girl.*F.Sg* had fallen.*M /* The woman saw that the necklace.*F.Sg* had fallen.*M*). They observed an N400 effect when the subject was animate, wherein conceptual information or semantic gender was at play. Additionally, a P600 was observed for both the conditions, which was attributed to reanalysis processes.

A similar study was conducted by Barber et al. (2004) who examined the comprehension of grammatical and semantic gender in Spanish. For their study they employed noun-adjective combinations in sentences, in which they manipulated the type of nouns (i.e. animate/inanimate) for the different types of gender, and the agreement (correct/mismatch). A LAN-P600 biphasic effect was observed irrespective of the animacy of the noun, whereby the LAN was interpreted as suggestive of the detection of a feature mismatch between the two participating elements. The P600 was ascribed to the fact that gender agreement was a part of the syntactic component during the comprehension process. This study thus did not find any differences in the comprehension of syntactic and semantic gender in Spanish. Recently Popov (2017) examined determiner-noun agreement in Italian, where the animacy of the noun was manipulated to denote semantic and syntactic gender. Further, the study had two different modalities as well: visual and auditory. With respect to the visual modality there were no differences whatsoever with regards to the differing animacy of the noun. A LAN-P600 pattern was observed for both the violation conditions. However, in the auditory modality, a difference ensued based on the differing animacy of the noun. When the noun was animate (semantic gender), a P600 was observed whereas when the noun was inanimate (syntactic gender), a N400-P600 pattern ensued. The N400 was attributed to the retrieval of the gender feature of the noun by lexical access from the lemma. The P600s on the other hand were said to be related to integration issues.

Thus from the studies cited above, it is clear that there is no consensus for the syntactic-semantic gender divide either with respect to the type of components observed or with respect to whether the processing mechanism for the two types of gender is similar or different. Interestingly, with the exception of Deustch and Bentin (2001) other studies have probed into modifier-head agreement relations thus leaving verb-agreement relations relatively unexplored from this vantage point.

#### 2.1.2 Processing gender agreement in Hindi

A few previous studies have investigated the comprehension mechanism of gender agreement in Hindi. One of the earliest ERP studies in the language, Nevins et al. (2007) made use of the future tense verb form (which inflects for all the three phi-features i.e. person, number and gender) to probe into the question of whether the different phi-features have similar or different processing mechanisms between the verb and the subject argument. Further, they also intended to examine whether 1-feature violations were qualitatively different from 2-feature violations. The study had the following critical conditions: (1) Gender agreement violations (2) Number agreement violation (3) gender+number agreement violation (4) number + person agreement violation. The results showed a P600 component being elicited irrespective of the phi-feature being violated. Further, there seemed to be no difference between the 1-feature and 2-feature conditions except for the person+number violations that showed an increase in the amplitude of the P600. The results seemed to point towards agreement being a unified process irrespective of the type and number of features processed. However, there are two interesting points to note about the results of the study. The first one is the absence of a LAN effect that has been found for subject verb agreement violations across languages. The second is the larger amplitude of the P600 in case of person+number violations. Regarding the absence of LAN, the authors argue that even in other studies the factors due to which LAN is present or absent are not very clear. Besides, they also make the point that in the earlier studies, observing a LAN was attributed to vary as a function of the auditory vs. visual presentation mode or as a function of the working memory capacity and not as a direct reflection of morphosyntactic errors. As for the larger P600 for person+number violations, it is said to be due to the special status of the person feature in linguistic theory in general and also in the language under study.

The ERP studies by Bornkessel-Schlesewsky et al. (2008) and Choudhary (2011) examined long distance agreement, a phenomenon in which the verb agrees not with its own argument but with an argument outside the clause i.e. the object of the embedded clause. The authors had two main aims in their study: the first was to investigate if the pattern found in case of LDA violations was the same as in case of agreement violations in simple clauses. The overarching objective here was to test whether the two kinds of agreement i.e. within clause and extra-clausal agreement had similar or different processing mechanisms. The other aim was to see if the case marking of the matrix subject had a role to play with regards to agreement dependencies. The following factors were manipulated in the experiment: case marking of the matrix subject (nominative/ergative); agreement properties of the infinitive (masculine/feminine); agreement properties of the control verb (masculine/feminine). The study comprised of three critical positions: (1) Case-marking on the first NP: only the highest ranked nominative case marked arguments agree with the verb, and LDA is possible with only when the subject is case marked with ergative or dative in Hindi. Thus, in the case of nominative marked subject the agreement with the control verb was highly predictable as compared to an ergative marked argument. (2) Agreement properties of the infinitive: LDA occurs only when the infinitive agrees with its O argument. The point here was to see how the infinitive influenced the agreement of the control verb. (3) Agreement properties of the control verb: here, a LAN-P600 or P600 effect was expected in line with previous studies. The ERPs at the control verb were analysed in terms of the following factors: case marking of the matrix subject (ergative vs. nominative) & gender agreement of control verb & infinitive: masculine-masculine (MM), feminine-feminine (FF), and masculine-feminine (MF). The results were a reflection of modulation of all the factors mentioned above. When the matrix subject was nominative case marked, in the MM condition a P300 was elicited whereas in the FF and MF condition an N400 was observed, with the effect being larger for the FF condition. The P300 was attributed to the fact that when the NP1 is nominative marked and masculine then there is a high expectation for agreement with the control verb. When this expectation is not fulfilled, a P300 ensued. The N400 in the other two conditions is said to be due to the agreement mismatch between the matrix subject and the control verb. Additionally, the FF condition also engendered a late positivity. In case of the ergative marked matrix subject, the FM condition showed an N400 effect that was ascribed to the fact that when the matrix subject is ergative case marked, there was an expectation for the agreement in the matrix clause to match that of the infinitive. Violation of this leads to the elicitation of an N400. In a nutshell, this study highlights the distinct processing mechanisms for LDA and agreement in simple clauses as evident from the different ERP components observed when the matrix subject is nominative (corresponding to agreement in simple clause) as opposed to when its ergative (corresponding to LDA).

From the studies described above, an important point that becomes clear is that gender agreement even in one and the same language has many facets depending upon various factors that are manipulated, one of them being the construction involved in the agreement relation. Furthermore, ERP effects elicited by violations of gender agreement in the language may differ qualitatively depending upon these factors.

### 2.2 Neurophysiology of Gender Retrieval

As in the case of gender agreement processing and the syntactic-semantic gender divide, studies focusing on gender retrieval on different languages have also revealed divergent results. Caffarra, Janssen & Barber (2014) examined potential evidence for the existence of the two-route model while comprehending article-noun pairs in Spanish. They manipulated the noun transparency as well as the agreement between the article-noun pair. Additionally, this study employed a visual half field method to investigate possible hemispheric distinctions in gender agreement processing. ERP effects were reported in two time windows: 350-500 ms & 500-750 ms. In the earlier time window, a negativity effect around 400 ms (interpreted as N400) was found in both the hemispheres. However, while the main effect of agreement was found in both hemispheres in this time window, the main effect of transparency could be observed only in the left hemisphere, whereby the ERPs for transparent nouns were more negative going as compared to opaque ones. In the latter time window, a main effect of agreement was present in both the hemispheres. A main effect of transparency was again observed only in the left hemisphere. However, in the right hemisphere significant interactions were seen between transparency and agreement suggesting different processing routines for transparent and opaque nouns. The fact that there was no interaction between agreement and transparency in the initial time window seems to suggest that at least in the earlier stages agreement computation is independent of the noun ending. The increased negativity for transparent nouns in the left hemisphere in the earlier time window was attributed to the fact that twin sources of lexical as well as form-based information are available while processing transparent nouns, but in case of opaque nouns only the lexical information is available. This increased source of information was conjectured to be the reason for the larger negativity for transparent nouns.

Caffarra, Siyanova-Chanturia, Pesciarelli, Vespignani and Cacciari (2015) investigated if the dual route mechanism worked in case of determiner-noun gender agreement violations in a sentential context in Italian. The study manipulated the transparency of the noun as well as the agreement between the determiner and the noun. Results revealed a main effect of transparency as well as a main effect of agreement. A frontal negativity for the transparent nouns was observed in the frontal region between 350-550 ms as well as 550-750 ms, and a positivity was found for the same between 750-950 ms in the posterior sites. A LAN ensued between 350-500 ms for the agreement mismatch conditions followed by a positivity between 750-950 ms. The authors posit that the agreement mismatch elicits a LAN-P600 effect. Also, the transparency effect is said to be suggestive of the fact that the parser is sensitive to the noun-ending cues and that these cues could be detected as early as 350 ms after the onset of the noun and remain active till as late as 950 ms. Additionally, the topographic differences in the negativity seen in case of the transparency effect (frontal) and the agreement effect (LAN) are said to be indicative of the fact that these two factors involve different neurocognitive mechanisms. This argument is bolstered by the fact that no interaction was seen between transparency and agreement. Simply put, although the parser detects noun-ending cues with respect to gender during processing, this does not affect the computation of agreement.

Caffarra and Barber (2015) studied determiner-noun agreement in Spanish in a sentential context to examine if the dual-route mechanism influenced gender agreement computation in the language. The authors manipulated the noun transparency as well as the agreement between the determiner and the noun in the experiment. They saw a two-fold result: a main effect of agreement as well as a main effect of transparency. However, no interaction was observed. For the agreement violation conditions, a LAN-P600 effect was observed whereby the LAN was said to be due to the detection of a grammatical agreement mismatch, and the P600 due to repair and reanalysis processes. Also, they saw a main effect of transparency between 200-500ms with the transparent nouns showing a greater negativity than opaque nouns at fronto-central electrode sites. The greater negativity seen for transparent nouns is attributed to the fact that for transparent nouns both the formal as well as lexical routes are activated whereas for the opaque nouns only the lexical route needs to be activated. Thus, in case of transparent nouns a greater amount of information needs to be activated and integrated as compared to opaque nouns, which is reflected in the increased negativity.

A point to note here is that the authors did not observe an interaction between the transparency and agreement effects which in essence means that though the parser is sensitive to gender-ending cues, the gender-ending cues do not affect agreement computations. The authors offer an explanation to this end that once the gender of a noun is retrieved (through either of the routes), the route through which the gender was actually retrieved does not make a difference in computing agreement dependencies.

Resende, Mota and Seuren (2018) analysed determiner-noun and noun-adjective agreement in Brazilian Portuguese while manipulating the noun transparency. The goal of the study was to see if noun transparency elicited different effects thus providing evidence for the dual-route model. Additionally, the authors also were interested to examine if determiner-noun and noun-adjective agreement involved similar or different processing mechanisms. A point to be noted is that the authors used only inanimate nouns in the study and the critical position in determiner-noun agreement conditions was the noun, while it was the adjective in noun-adjective agreement conditions. Determiner-noun as well as noun-adjective agreement violations elicited a LAN-P600 effect in the 300-500 ms and 500-900 ms time-windows respectively. This effect did not differ with respect to the noun transparency. Also, they observed a positivity in the 500-900 ms for the noun-adjective condition, between regular and irregular nouns. This is said to be due to the fact that the mismatch between a regular noun and an adjective is more easily detected than between an irregular noun and an adjective due to frequency differences between the two. In sum, the results observed in this study do not support the claims for the presence of a dual-route model in the language.

Thus, cross-linguistic data pertaining to the investigation of the dual-route mechanism are inconclusive. While some languages confirm the presence of the dual-route mechanism others do not. However, a pattern that emerges from these studies is that even in languages that show the presence of the dual-route mechanism, the gender retrieval mechanism does not seem to affect the comprehension of agreement relations. But the point to be noted here is that in these studies the critical point is the noun itself, or in some cases the adjective, but not the verb.

## 3. The Current Study

The primary goal of the current study was to investigate gender agreement computations in Hindi, a verb-final Indo-Aryan language. With respect to verb-argument agreement relations, the gender feature is realized on the noun (target) as well as the main or the auxiliary verb depending upon the tense in use in the sentence.

Theoretically, the agreement dependencies in the language are not dependent on the intrinsic properties of the noun phrase involved^4^. The present study however manipulated the animacy of the subject argument participating in the agreement relation to investigate whether the argument’s features such as animacy have a bearing in processing agreement dependencies. In the psycholinguistic literature, most works have examined animacy with respect to its role in thematic hierarchizing in transitive constructions (Roehm, Schlesewsky, Bornkessel, Frisch and Haider, 2004), syntactic ambiguity resolution (Philipp, Bornkessel-Schlesewsky, Bisang, Schlesewsky, 2008) or form-to-meaning mapping (Bornkessel and Schlesewsky, 2009b). As discussed earlier, in the domain of agreement comprehension, animacy comes into play in studies that examine the processing differences between semantic and syntactic gender. In this regard, most of these studies have taken into consideration modifier-head agreement with the exception of Deutsch & Bentin (2001) who examined the processing of agreement between a predicate and its argument in Hebrew. Likewise, the results of these studies have differed based on the language investigated and structures involved in agreement. Thus from an experimental viewpoint there is a dearth of data with regard to how gender agreement relations would be computed similarly or differently between an animate subject noun and its verb as opposed to an inanimate subject noun and its verb in an intransitive construction.

The present study strives to fill this research gap. Hindi presents an opportunity to investigate this while additionally allowing to examine the differences between processing of semantic and syntactic gender in the same sentences. In most previous studies on animacy, a transitive structure was used whereby the participants could use the animacy information at the second NP to compute relational information between the two arguments in the sentence. In this study, the use of intransitive structures implies that at the position of the verb both the animacy of the NP1 and the agreement information will have to be amalgamated thus informing whether or not animacy plays a role in agreement computation in the language. Based on these ideas and on the studies described in sections 2.1 and 2.2, we hypothesize that if the animacy of the subject noun influences the computation of subject-verb agreement relations, we expect qualitatively different ERP components between the animate subject and the inanimate subject conditions in the study. This could also be an indicator that the way the gender retrieval is performed influences gender agreement computation, and animacy could emerge as one of the factors that impacts gender retrieval of nouns in the language. It could also be argued that this reflects the ability of the human parser to distinguish between semantic and syntactic gender in the language. Alternatively, if the parser glosses over the animacy of the subject noun, similar to that in Barber et al. (2004), qualitatively similar ERP components should be elicited suggesting similar mechanisms behind syntactic and semantic gender processing in the language. Based on the previous studies on agreement in Hindi (i.e. Nevins et al., 2007; Bornkessel-Schlesewsky et al., 2008 and Choudhary, 2011), we expect either a negativity-positivity pattern or a sole positivity for the two conditions or a negativity-positivity in case of animate subject arguments and negativity in case of inanimate subjects. However, due to the fact that heterogeneous ERP effects have been reported not only across languages but also within languages, the exact ERP components remain to be seen. Besides, this would also specify that gender retrieval and computation are independent processes without any influence on each other.

## 4. Materials and Methods

### 4.1 Participants

Twenty-two native speakers of Hindi (mean age = 20.05; age range = 18-30), mostly students and staff at the Indian Institute of Technology Ropar, living in Ropar, India participated in the experiment. Before the start of the experiment, the participants were briefed about the experiment and informed consent was obtained from them for the use of their data for academic purposes. Also, they were given suitable remuneration for their time. The research protocol for the experiment was approved by the Institutional Ethics Committee (Human) of the Indian Institute of Technology Ropar. All the participants were right-handed as evaluated by an adapted version of the Edinburgh Handedness Inventory (Oldfield, 1971) in Hindi. The participants had normal or corrected to normal vision. Also, they had no known neurological disorder at the time when they participated in the experiment. All the participants spoke English in addition to Hindi and reported to having learnt Hindi before the age of six. Further, some of them also spoke more language(s) in addition to these two languages. The participants of the experiment hailed predominantly from the Hindi-speaking states of Delhi, Uttar Pradesh or Madhya Pradesh in India.

### 4.2 Materials

#### 4.2.1 Critical Conditions

The experiment comprised of four intransitive sentences in an item set. 72 such sets were created for the purpose of the experiment. The four conditions differed on the basis of gender agreement violation and the type of the subject argument. The factors and levels with appropriate codes for each condition are shown below.

The CA condition exhibits agreement with the animate subject and the CI condition exhibits agreement with the inanimate subject, both of them grammatical. In the MA condition, there is a gender agreement mismatch with the animate subject argument, and therefore it is ungrammatical. Similarly, in the MI condition, there is a gender agreement mismatch with the inanimate subject argument and therefore it is ungrammatical. The subject argument in each condition is a unique animate/inanimate common noun that was marked with nominative case. All the arguments used in this experiment were masculine. The verbs used in the set from 1-36 were unique and they were repeated from the set 37-72. In all the conditions the verb was in imperfective aspect. The stimuli were controlled for length. Table 2 provides a sample set of the critical conditions.

**Table 1:** Factors and Levels.

| FACTOR | LEVEL | CONDITIONS |
| --- | --- | --- |
| Agreement | C: Correct | CA, CI<br>MA, MI |
|  | M: Mismatch |  |
| Animacy of the subject | A: Animate |  |
|  | I: Inanimate |  |

**Table 2:** Experimental test stimulus^5^.

| CONDITION | SAMPLE |  |  |
| --- | --- | --- | --- |
| **CA** | किसान<br>**Kisaan**<br>Farmer.3SG.M.NOM | गिरता<br>**gir-taa**<br>fall-IPFV.3SG.M | है<br>**hai**<br>Aux |
|  | “The farmer falls (down).” |  |  |
| **\*MA** | किसान<br>**Kisaan**<br>Farmer.3SG.M.NOM | गिरती<br>**gir-tii**<br>fall-IPFV.3SG.F | है<br>**hai**<br>Aux |
|  | “The farmer falls (down).” |  |  |
| **CI** | परदा<br>**Pardaa**<br>Curtain.SG.M.NOM | गिरता<br>**gir-taa**<br>fall-IPFV.SG.M | है<br>**hai**<br>Aux |
|  | “The curtain falls (down).” |  |  |
| **\*MI** | परदा<br>**Pardaa**<br>Curtain.SG.M.NOM | गिरती<br>**gir-tii**<br>fall-IPFV.SG.F | है<br>**hai**<br>Aux |
|  | “The curtain falls (down).” |  |  |

#### 4.2.2 Fillers

Filler sentences were employed to introduce diverse structures in the stimuli so as to avoid strategic responses from participants. Twenty-four fillers were constructed each in six types, thus resulting in a total of 144 filler sentences. These were interspersed with the critical sentences and pseudo randomised before presentation. The nouns used in the filler sentences were different from those in the critical sentences.

#### 4.2.3 Item Distribution

The 288 experimental sentences were divided into two lists, each comprising of 144 sentences. Each set of 144 critical items was interspersed with the 144 filler sentences and pseudo randomised as described above. The 144 critical sentences along with 144 fillers made up one session. There were two sessions in the experiment. Each of these sessions were counterbalanced across participants and randomized.

#### 4.2.4 Acceptability Judgment

Participants performed an acceptability judgment task after the stimulus was displayed, whereby they had to press a green button in the response pad if they found the stimulus displayed earlier to be acceptable, or alternatively a red button if they found the stimulus to be unacceptable.

#### 4.2.5 Comprehension Question

To make sure that the participants were attentive while the stimulus was being displayed, a comprehension question followed each sentence. Comprehension questions that required a ‘yes’ response were essentially the stimulus sentence along with the question particle “kya” in the beginning. Half of the comprehension questions were correct while the other half was incorrect (either the first argument or the verb was changed with respect to the stimulus displayed earlier). There were equal number of comprehension questions in which either the verbs or the first argument was changed.

### 4.3 Method

#### 4.3.1 Procedure

The experiment was conducted at the Language and Cognition Lab, Indian Institute of Technology Ropar. It consisted of a practice session followed by the actual experiment, with all the activities including electrode preparation and stimulus presentation lasting approximately two hours.

Adult right-handed native speakers of Hindi participated in the experiment. As soon as they arrived at the lab, the procedure and the tasks to be performed during the experiment were explained to the participants. A printed instruction sheet was provided. However, the question under investigation in the experiment was not revealed to them so that unbiased data could be acquired. Then they filled a consent form giving an informed consent for their participation. They then filled a Hindi version of the Edinburg handedness questionnaire to evaluate their handedness.

Then, the head measurements of the participants were taken and the Hydrocel GSN net was placed on their scalp. After this, they were taken to a soundproof chamber, in which they were seated on a comfortable chair at a certain distance (1 meter) from the screen (20 inch LCD) on which the stimulus would be presented. Further, they were briefed again to avoid any drastic movements and especially eye blinks during the presentation of the stimulus. After this, they were given a practice session to familiarize themselves with the experimental trial structure. After this the actual experiment commenced. After each block of 48 trials, a short break was provided to the participants. At the end of the experiment, participants were requested to fill a questionnaire regarding their experience about the experiment.

#### 4.3.2 Practice Session

The Practice session consisted of 10 trials presented visually. The manner of display was identical to that of the actual experiment so as to give the participants a feeling of the actual experiment. The sentences were presented visually in a word-by-word fashion (with case markers appearing together with the word) at the center of the presentation monitor screen. As soon as a trial began, the participants would see a ‘+’ fixation sign at the center of the screen for a period of 1000ms. A blank screen for a duration of 200 ms followed. Each of the words were presented for a period of 650 ms followed by an inter stimulus interval (ISI) for a duration of 200 ms. After the fixation an adverb appeared on screen followed by the NP1, verb and the auxiliary. As a cue to the acceptability judgment task a “???” sign appeared on the screen just after the auxiliary. Here, the participants had to judge whether the stimulus they just saw was acceptable or not. Accordingly, if they found it to be acceptable, they had to press the green button on the response pad. If they found the stimulus to be unacceptable, the participants had to press the red button. After the participants answered or after a maximum duration of 1500 ms, whichever is earlier, a comprehension question pertaining to the stimulus they saw appeared on the screen. Here again, the participants had to press the green button on the response pad if their answer was “yes” or the red button if their answer was “no”. The maximum time duration for this task was 3000 ms. Half of the comprehension questions were correct and half were incorrect. In incorrect comprehension questions, the NP1 or the verb was changed. A schematic diagram of the presentation schema is shown in Figure 1.

**Figure 1:**
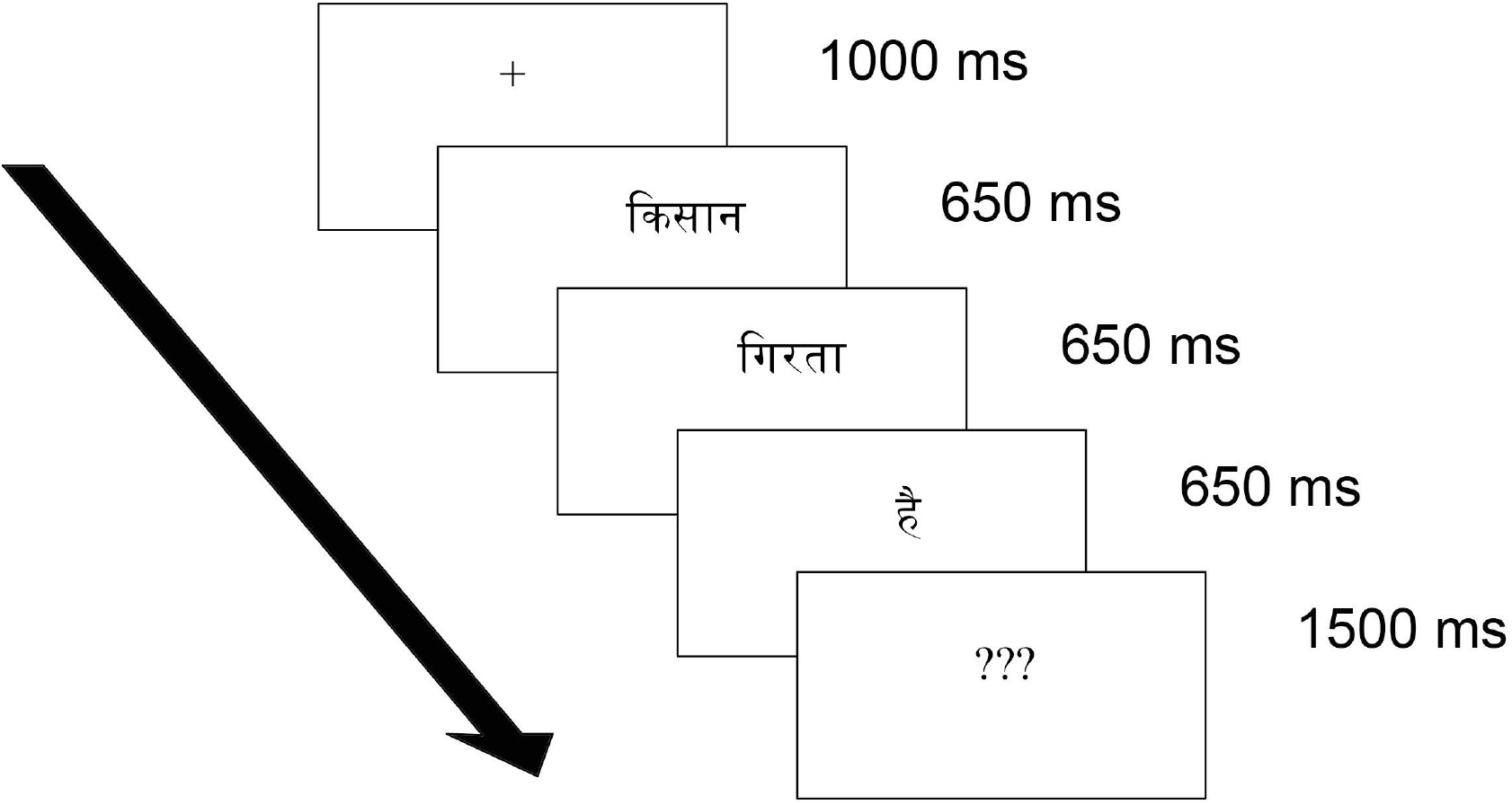
Stimulus presentation Schema.

Participants were requested to avoid blinks during the presentation of the words but they could blink while performing the two tasks.

As soon as the practice session ended, a break was provided to the participants during which they were asked if they were fine with going ahead and performing the experiment. After this the actual experiment began.

#### 4.3.3 Experimental Session

The experimental session consisted of 288 sentences divided into 6 blocks of 48 sentences each. The stimulus presentation was in the same manner as described in the practice session. The participants were given a short break after the completion of each block.

#### 4.3.4 EEG Data Acquisition and processing

Scalp activity was recorded by means of 129 AgAgCl electrodes (128+VREF) fixed at the scalp by means of a Hydrocel Geodesic Sensor Net 128 channel (Philips Neuro, OR, Eugene). The vertex electrode served as the reference electrode. However, the recordings were re-referenced offline to the average of the two mastoids (Molinaro, Barber and Carrerias, 2011). For monitoring EOG (Electro-oculogram) data, the electrodes were placed as follows: for capturing horizontal eye movements two electrodes were placed at the outer canthus of each eye (electrode numbers 125, 128) and for vertical eye movements there were a total of 6 electrodes, with four placed above the eye (8, 14, 21, 26) and two below (126, 127). The interelectrode impedance was kept below 50 kΩ as per system recommendations. All EEG and EOG data was amplified using a Net Amps 400 Amplifier. The data was recorded at a sampling rate of 500 Hz.

The EEG data was analysed using EEGLAB Toolbox (Delorme & Makeig, 2004). The acquired raw EEG data was filtered using a bandpass filter of 0.3-30 Hz followed by offline re-referencing to the average of the right and left mastoids. Thereafter, automatic bad channel rejection procedure was performed using the ‘pop_rejchan’ function in the EEGLAB toolbox. The resulting data was again filtered at 1 Hz as per EEGLAB recommendations before an extended infomax ICA (Iriarte, Urrestarazu, Valencia, Alegre, Malanda, Viteri, Artieda, 2003) was applied on the continuous EEG data. SASICA (Chaumon, Bishop and Busch, 2015) was used to identify and reject the artifact laden ICA components. Subsequently, the ICA weights were transferred to the pre-1Hz data followed by interpolation. Then the data were epoched for 1200ms after the onset of the critical word (the verb) relative to a 200m pre-stimulus time after which the data were averaged per subject per condition. Finally, the grand average was computed based on 22 participants.

#### 4.3.5 EEG Analysis

Separate analyses were conducted for midline and lateral regions. For the statistical analysis, mean value of a group of electrodes comprising of one region of interest (ROI) were extracted using the ERP measurement tool of the EEGLAB toolbox. As such, the midline regions were divided into three ROIs: middle frontal (18, 16, 10, 19, 11, 4, 12, 5), middle central (13, 6, 112, 7, 106, 31, 80, 54, 55, 79) and middle posterior (61, 62, 78, 67, 72, 71, 77, 76). The lateral regions had six ROIs: left frontal (23, 24, 20, 27, 28, 29, 34, 35), left central (40, 41, 42, 36, 37, 30, 46, 47), left posterior (51, 52, 53, 58, 59, 60, 65, 66), right frontal (3, 123, 124, 117, 118, 110, 111, 116), right central (98, 93, 87, 102, 103, 104, 105, 109) and right posterior (86, 92, 97, 96, 90, 91, 84, 85). The time windows for which statistical analyses were performed were based on visual inspection of the data.

The repeated measures ANOVA had the following within subject factors: 1. Agreement (2 levels: correct and mismatch) 2. Animacy (2 levels: animate and inanimate) 3. Anteriority (3 levels: frontal, central and posterior). In case of the ANOVA for lateral regions it included an additional factor of Hemisphere (2 levels: right and left). Significance levels were kept at *p* <0.05. For avoiding sphericity violations, Huynh and Feldt-correction was applied to the *p-*values (Huynh and Feldt, 1970).

## 5. Results

### 5.1 Behavioural Data

The answering accuracy and the mean reaction time for the acceptability judgment as well as the comprehension questions were first extracted using the E-data aid application of E-Prime 2.0 (Schneider, Eschman, & Zuccolotto, 2002). Further analysis was done using R (R Core Team, 2017). Table 2 shows the answering accuracy and the mean reaction times for both the tasks in the experiment. Further statistical analysis of the behavioural data was done using repeated measures ANOVA(s) that had within-subject factors agreement and animacy and the random factors participants (F1) and items (F2).

The behavioural results as depicted in the Table 3 (i.e. the mean acceptability for the acceptability judgment task) above show that the ungrammatical conditions were rated as significantly less acceptable as compared to their grammatical counterparts. Further, the reaction time for this task also reveals longer reading times for the two ungrammatical conditions in comparison to the correct ones thus suggesting a slowing down whenever a violation was encountered.

**Table 3:** Mean Acceptability and Accuracy rates along with reaction times for the two tasks in the experiment. Standard deviations are given in parenthesis.

| Condition | Acceptability Judgment |  | Comprehension Question |  |
| --- | --- | --- | --- | --- |
|  | Mean Acceptability (in %) | Reaction Time (in ms) | Mean Accuracy (in %) | Reaction Time (in ms) |
| CA | 91(15) | 486(183) | 96(03) | 962(232) |
| *MA | 30(27) | 510(191) | 94(06) | 1067(292) |
| CI | 87(16) | 491(190) | 97(04) | 1004(269) |
| *MI | 37(22) | 505(186) | 92(08) | 1037(298) |

A Repeated measures ANOVA(s) for the acceptability judgment task revealed a main effect of AGREEMENT (*F*_*1*_(1,21) = 69.6, *p*<0.001 ; *F*_*2*_(1,71) = 480.3, *p*< 0.001). The ANOVA for the reaction time of the acceptability judgment task reached significance only in the analysis by items showing a main effect of AGREEMENT (*F*_*2*_(1,71) = 5.89, *p*< 0.0177).

The repeated measures ANOVA for the comprehension question task showed a main effect of AGREEMENT (*F*_*1*_(1,21) = 6.77, *p*< 0.0166; *F*_*2*_(1,71) = 7.88, *p*< 0.00644). Statistical analysis of the reaction time for the comprehension question revealed a main effect of AGREEMENT (*F*_*1*_(1,21) = 15.61, *p*< 0. 00073; *F*_*2*_(1,71) = 30.82, *p*<0.001) as well as an interaction of ANIMACY x AGREEMENT (*F*_*1*_(1,21) = 15.9, *p*< 0.00067; *F*_*2*_(1,71) = 12.3, *p*< 0.000788). When the interaction between ANIMACY x AGREEMENT was resolved for ANIMACY, it revealed a main effect of AGREEMENT for the animate condition (*F*_*1*_(1,21) = 21.52, *p*<0.0001; *F*_*2*_(1,71) = 37.99, *p*<0.001) and only in the analysis by items for the inanimate condition (*F*_*2*_(1,71) = 5.59, *p*<0.02).

### 5.2 ERP Data

ERPs were calculated for each participant from 200 ms before the onset of the verb till 1000 ms afterwards (−200 ms to 1000ms). For the statistical analysis of the ERP data, repeated measures ANOVA(s) were computed with the within participant factors AGREEMENT and ANIMACY for mean amplitude values per time-window per condition in the six lateral and three midline ROIs (Regions of Interest) as described in Section 4.3.5. The time windows were selected based on visual inspection of the data. Visual inspection of the grand-averaged data revealed a widespread negativity effect for the inanimate violation condition as opposed to its acceptable counterpart between 500-900 ms (Figure 2), and a positivity effect for the animate violation condition as opposed to its acceptable counterpart between 650-850 ms (Figure 3).

**Figure 2:**
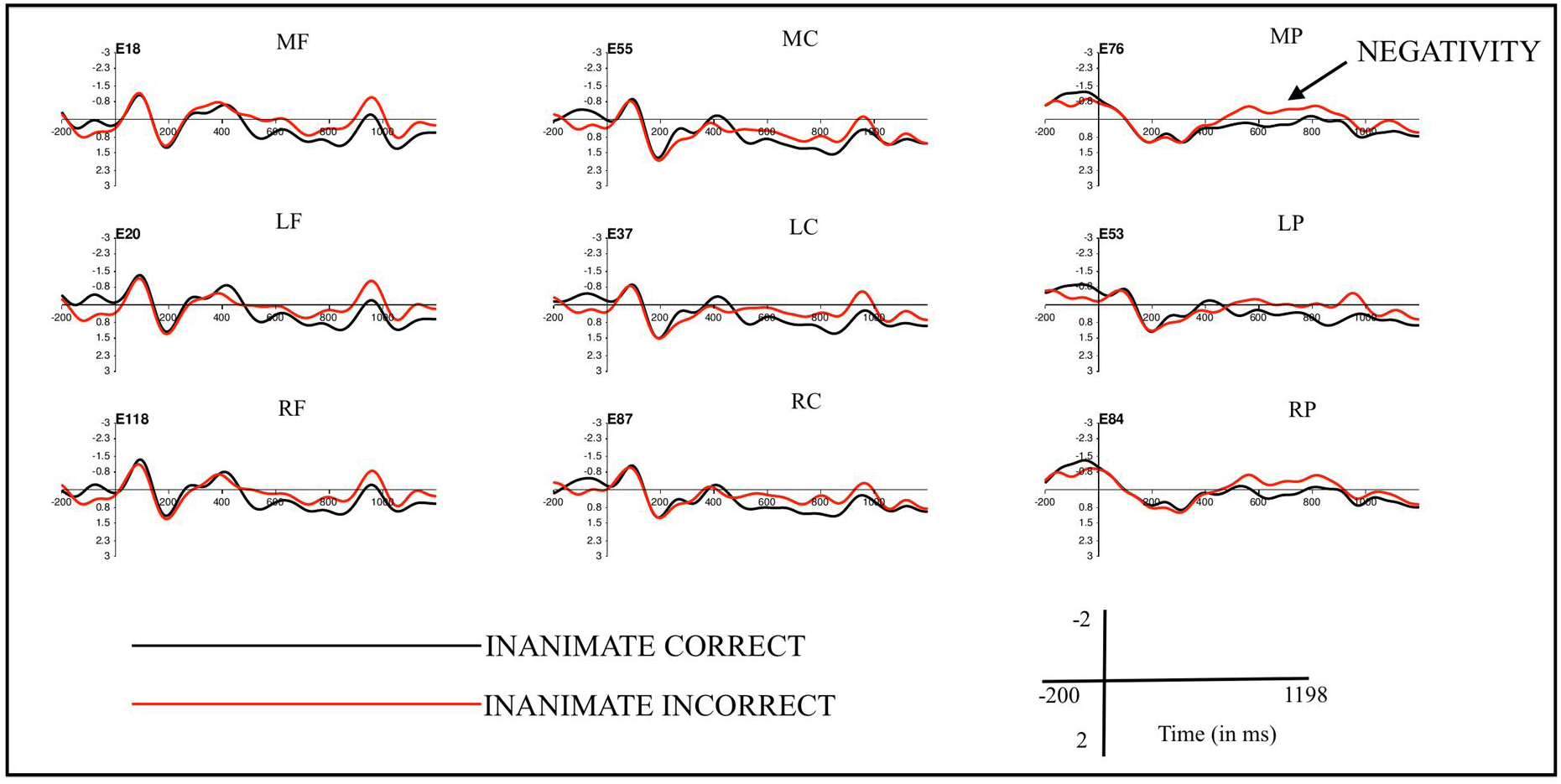
Grand averaged ERPs at the position of VP (n = 22) comparing the correct and incorrect inanimate conditions. The black line is for the correct inanimate condition and the red line for incorrect inanimate condition. Abbreviations: MF: Middle frontal, MC: Middle central, MP: Middle posterior, LF: Left frontal, LC: Left central, LP: Left posterior, RF: Right frontal, RC: Right central, RP: Right posterior

**Figure 3:**
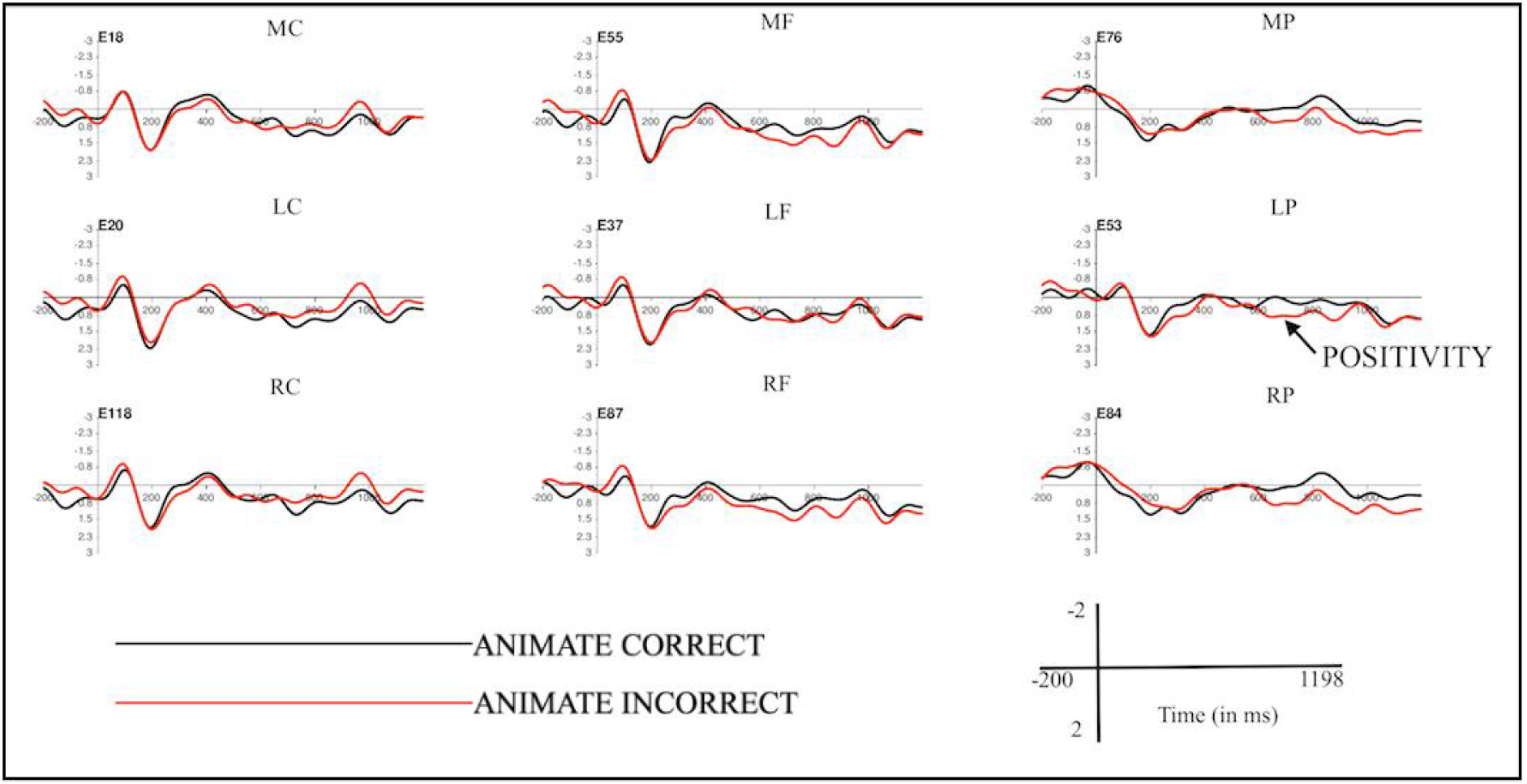
Grand averaged ERPs at the position of VP (n = 22) comparing the correct and incorrect animate conditions. The black line is for the correct animate condition and the red line for incorrect animate condition. Abbreviations: MF: Middle frontal, MC: Middle central, MP: Middle posterior, LF: Left frontal, LC: Left central, LP: Left posterior, RF: Right frontal, RC: Right central, RP: Right posterior

#### VP: (Verb)

##### Time Window: 500-900 ms

The analysis for this time window revealed an interaction of ANIMACY x AGREEMENT (midline: *F*(1,21) = 10.86, p<0.003; lateral: *F*(1,21) = 8.38, p<0.008). Resolving this interaction by ANIMACY, revealed a main effect of AGREEMENT for the inanimate condition (midline: *F* (1,21) = 7.52, p<0.012; lateral: *F* (1,21) = 6.30, p<0.020).

##### Time Window: 650-850 ms

The analysis for this time window revealed an interaction of ROI x ANIMACY x AGREEMENT (midline: *F*(2,42) = 3.65, p<0.052 ; lateral: *F*(5,105) = 3.30, p<0.032). Resolving this interaction by ROI revealed an interaction of ANIMACY x AGREEMENT in the middle central (*F* (1,21) = 5.39, p< 0.030) and Middle posterior (*F* (1,21) = 21.15, p<0.001) regions for the midline sites and left posterior (*F* (1,21) = 13.01, p< 0.001) right central (*F* (1,21) = 7.42 p< 0.012) and right posterior region (*F* (1,21) = 11.58, p< 0.002) for the lateral sites.

When the interaction between ANIMACY x AGREEMENT was resolved for ANIMACY, it revealed a main effect of AGREEMENT for the animate condition at the middle posterior region (*F* (1,21) = 6.24, p< 0.020) and for the inanimate condition at the middle central (*F* (1,21) = 4.15, p< 0.054) and the middle posterior region (*F* (1,21) = 5.50, p< 0.028).

Further for the lateral sites it revealed a main effect of AGREEMENT for the animate condition at the left posterior region (*F* (1,21) = 5.52, p< 0.028) and for the inanimate condition at the right posterior region (*F* (1,21) = 4.66, p< 0.042).

### 5.3 Summary of Results

To sum up, the statistical analysis supports the presence of a negativity effect for the inanimate violation condition and a positivity effect for the animate violation condition. Further, the statistics also showed a significant interaction of animacy and agreement both for the animate and the inanimate conditions between 650-850ms. However, the significance here for the inanimate condition is due to the overlapping time-windows of the negativity in the 500-900 ms and 650-850 ms time-windows. There is no evidence for a positivity effect for the inanimate condition as illustrated by Figure 2.

## 6. Discussion

The present study investigated the neural basis of semantic and syntactic gender agreement in verb-argument relations in Hindi. We hypothesized that, either the processing of semantic and syntactic gender will be evidenced by similar ERP components (Hypothesis 1) or alternatively, they will involve different comprehension mechanisms (Hypothesis 2). Our results support Hypothesis 2, with the syntactic gender condition (inanimate subjects) engendering a negativity effect and the semantic gender condition (animate subjects) eliciting a late-positivity effect. The fact that we did not find a late-positivity effect for the syntactic gender condition is in contrast to results from earlier studies on agreement processing, which have either reported a biphasic effect or a P600 effect.

Our results appear to indicate that the underlying mechanisms behind the processing of the two kinds of gender agreement relations are different in Hindi. The fact that the agreement is with an animate subject in case of semantic gender whereas it is with an inanimate subject in case of syntactic gender is suggestive of the importance of the role of animacy in computing the two kinds of gender agreement. Thus, the two types gender agreement differentiated by the animacy of the noun in Hindi provide counter-evidence to the view that gender agreement is a monolithic phenomenon indifferent to finer distinctions.

A point to highlight here is that the noun has been the critical focus of studies related to semantic and syntactic gender processing and those that discuss different routes for gender retrieval. The current study however takes a different approach and looks at gender agreement relations from the vantage point of the verb. In this regard, the results of this experiment contribute crucial insights to our understanding of gender agreement. Also, the N400 reported in studies on gender agreement till date have been mostly associated with gender agreement violations involving animate nouns. An exception to this pattern was the study on Italian by Popov (2017), who observed an N400 when agreement with an inanimate noun was violated. The author argued that gender was stored as a feature of the lemma in Italian, and that full lexical access was required to retrieve its value during the comprehension of the noun, which is then said to give rise to the N400 observed in that study. Along similar lines, Caffarra and Barber (2015) did not observe an interaction of transparency and agreement in their study, thus concluding that gender retrieval and agreement computation were independent processes. The interaction of agreement and animacy observed in the current experiment suggest that gender retrieval and gender agreement computation are not independent of each other in Hindi. Besides, it also reinforces the argument that syntactic and semantic gender agreement in the language have separate mechanisms as evidenced by the distinct ERP components. Considering the arguments made in this section, we try to give a detailed functional interpretation of the ERP components observed in this study in the upcoming sections.

### 6.1 The negativity & the positivity

Traditionally associated with semantic processing (Kutas and Hillyard, 1980), the N400 component has also been reported in several existing studies investigating gender agreement (Barber and Carrerias, 2005; Wicha et al., 2004; Guajardo and Wicha, 2014; Deutsch and Bentin, 2001; Bornkessel-Schlesewsky et al., 2008; Choudhary, 2011). Gender retrieval may be accomplished by two routes: the lexical route and the computational or formal route. While the former route is available for retrieving the gender of any noun by lexical access, the latter is available only for those nouns that have certain distributional cues to indicate the gender of the noun. In this experiment, we manipulated the animacy of the noun to indicate the two genders: i.e. the syntactic and the semantic genders. If a noun was animate (human) in nature, its biological sex had a correspondence with its grammatical gender, whereas there was no such correspondence for inanimate nouns. This information about the relation between animacy and gender is readily available to native speakers of Hindi. Thus, the animacy information provided by our critical nouns would have been a salient cue in case of animate nouns such that, the gender of an animate noun would be readily transparent to our participants, whereas in case of inanimate nouns, the gender information would have remained opaque. Therefore, in order to retrieve the gender of the inanimate noun, the parser would have had no additional cues available that would then make the computational route accessible to it. The only option left to it would be to retrieve the gender information through the lexical route by full lexical access. This is reflected in the N400 effect observed in case of the syntactic gender (inanimate) violation condition, in which the agreement relation between the verb and the inanimate subject noun breaks down. Indeed, as Choudhary, Schlesewsky, Roehm & Bornkessel-Schlesewsky (2009) have argued, an N400 is elicited if an interpretively relevant cue is violated. On the other hand, in case the noun in the sentence was animate, the computational route would be open to the parser because the animacy information would transparently indicate the gender of the noun. The P600 effect elicited by violations in case of animate nouns seems to demonstrate that a different route is involved in the computation of the gender of animate nouns as compared to that of inanimate nouns.

It is worth noting that the latency of the negativity effect elicited by the syntactic gender condition (500-900ms) is quite different from the latency of the negativities observed for agreement violations in previous studies. The latency of negativities reported earlier usually tend to be in the time range of 300-600 ms. However, there are a few studies from varied domains of language processing, which have reported negativities in the time-range of 500-900 ms. These include the study by Popov (2017) mentioned earlier, who reported a negativity for the processing of syntactic gender, a study by Wittenberg, Paczynski, Wiese, Jackendoff and Kuperberg (2014) on the processing of light verb constructions in English, another by Coulson and Kutas (2001) on the processing of jokes, and a study by Huang, Lee and Federmeier (2010) on the processing of concrete concepts^6^.

As explained in the earlier sections, syntactic gender as such is arbitrary and opaque in comparison to semantic gender in Hindi, which leads to more resources being needed for the comprehension of syntactic gender agreement. Adding to the arbitrariness of syntactic gender in Hindi, a number of dialects of the language exist, which show variation in assigning gender to inanimate nouns. That is, the gender of inanimate nouns varies across different dialects of Hindi (based on discussions with native speakers).

Such complexities seem to be borne in mind by the native speakers resulting in the shifting of the latency of the negativity as compared to earlier studies. Converging support for taking dialectal variation into account and a processing complexity-based interpretation comes from a study on Arabic noun-adjective agreement involving animacy (humanness, to be specific) by Idrissi, Mustafawi, Khwaileh and Muralikrishnan (2019, pre-print of article in review). This issue does not arise in case of natural or semantic gender since it is linked to the biological sex of the referent. The explanation offered here for the latency shift of the ERP effect with regards to dialectal complexities and requirement of more resources for comprehension is in consonance with the account offered for processing complexities by Bornkessel-Schlesewsky & Schlesewsky (2019).

Another front where the dialectal complexities play an important part is the absence of positivity in the syntactic gender condition. The results point towards the fact that the native speakers keep in mind the dialectal variation of the syntactic gender while deciding the well formedness of the sentence as evidenced by the absence of a positivity in this particular violation condition. In this respect the results of the current study seem to mirror Choudhary (2011).

Apart from the negativity, the other effect we see is the positivity in case of animate subject violation. Such effects have been often observed in case of gender agreement mismatches. For the most part, the elicitation of a P600 is viewed from the perspective that gender agreement is a part of syntactic processing (Osterhout & Mobley, 1995; Osterhout, 1997; Hagoort and Brown, 1999; Hagoort, 2003; Martin-Loeches et al., 2006). A point to be noted here is that in most of these studies the authors compare the gender agreement condition to a semantic violation condition and make their inferences as such. The P600 component is also associated with repair and reanalysis processes in many studies of gender agreement (Barber and Carrerias, 2005; Barber et al., 2004; Popov, 2017) and more generally with conflict-monitoring processes (van de Meerendonk, Kolk, Chwilla, & Vissers, 2009). Another possible interpretation of P600 is its association with well formedness (Choudhary, 2011). The earlier Hindi studies on gender agreement (i.e. Nevins et al., 2007; Bornkessel-Schlesewsky et al., 2008 and Choudhary, 2011) have also reported a P600.

Nevins et al. (2007) observed a P600 for the gender mismatch between an animate subject and the verb, which is in line with results from our study for the animate subject-verb mismatch. The explanation we provide here can also account for the effect seen by Nevins et al. (2007). Choudhary (2011) reported an N400-P600 effect for the simple agreement condition and a N400 in case of long-distance agreement (LDA). Their results can be partly accounted for by the explanation offered for the results of our study. If we consider their LDA violation, there the agreement breaks down between the control verb and the object of the embedded clause, which is an inanimate argument (e.g. bicycle). This engendered an N400, which is in line with our own observation of an N400 in the present study when the agreement between an inanimate subject and the verb breaks down. By contrast, in case of the simple agreement condition, when the matrix subject is nominative-case marked, the violation occurs when the agreement breaks down between the control verb and the matrix subject that is animate (i.e. Ram). This violation is signalled by a biphasic N400-P600 effect that is different from what we have seen in our study when the agreement between an animate subject and the verb is violated. A possible speculation with regards to the cause of such divergent results in these two studies within the same language could be the fact that the type of construction used in the two studies were different: they used transitive constructions whereas our studies used intransitive sentences. However, this is by no means conclusive and is an issue for further research.

## 7. Conclusion and future directions

The ERP results of the current study (i.e. a positivity for animate subject-verb gender agreement violations and a negativity for inanimate subject-verb gender agreement violations) provides important insights regarding various facets of gender agreement computation in general, and with respect to Hindi in particular.

On a broader level, the fact that different comprehension mechanisms seem to be at play as evidenced by the different ERP correlates obtained for the animate and inanimate conditions is suggestive of the key role of animacy in gender agreement computation. Though theoretical accounts do not take animacy into consideration while deciding agreement patterns, electrophysiological results seem to speak otherwise. Apart from the current study, similar findings of animacy influencing gender agreement computation has been observed in Italian (Popov, 2017), as well as in Arabic (pre-print of Idrissi, Mustafawi, Khwaileh, & Muralikrishnan, 2019, article in review). These results establish animacy to be a key component at the heart of gender agreement comprehension across typologically different languages.

Specifically with respect to Hindi, the diversity of results from studies on gender agreement in the language indicate that gender agreement is not a monolithic phenomenon but is comprised of finer distinctions at multiple levels. The current study found differences in the processing of syntactic and semantic gender, which are in turn tied to the animacy of the subject noun. The differences in findings between the current study and those by Bornkessel-Schlesewsky et al. (2008) and Choudhary (2011) are indicative of the fact that different structures may engender different effects in the processing of a particular phenomenon within a language as detailed in section 6.1. These within-language results also highlight the fact that more research is needed to gain further insights into this domain. In this context, future work in the field can concentrate on manipulating the animacy of other arguments in a sentence (i.e. the object argument) with respect to gender agreement to see if comparable effects as seen in this study is observed. Additionally, since case plays an important role in determining agreement in the language, one can also explore how the workings of case and animacy are tied to the question of comprehending gender agreement in Hindi. Lastly, for all these different threads of research described above comparable studies can be done across the different dialects of Hindi.

## Footnotes

^1^ A noun is said to have semantic or natural gender if there is a correlation between its biological sex and its grammatical gender. Syntactic gender on the other hand entails no correspondence between the biological sex and the grammatical gender. In Hindi, human nouns and most non-human animate nouns have semantic gender whereas inanimate nouns mostly have syntactic gender.

^2^ Certain researchers have found evidence for only one route for gender retrieval (Levelt et al.1999) but the other group of researchers (Gollan and Frost, 2001) report evidence for two routes for gender retrieval. The first group talks about grammatical gender as an abstract entity that is stored in the mental lexicon and retrievable through lexical access. For those proposing two routes, the distinction between the two routes is made on the basis of the transparency of the noun. The 1^st^ route here is that of recovering gender through lexical access, which is available to all sorts of nouns. The second route is exclusively available to transparent nouns and is the computational route.

^3^ Though LAN has been mostly associated with detection of a morphosyntactic violation such as agreement, yet its stability in the recent past with respect to its elicitation in studies examining agreement violations has been the subject of debate in the literature (Molinaro, Barber and Carrerias, 2011;Tanner and Van Hell, 2014; Molinaro, Barber, Caffarra and Carrerias, 2015 and Tanner, 2015).

^4^ Theoretically, the properties of the noun phrases involved in an agreement relation except its case do not influence the agreement pattern. Case and agreement are symbiotic in Hindi with the rule being that a finite verb agrees with the highest non-overt case marked argument.

^5^ The following abbreviations are used in this study: M = Masculine, F = Feminine, SG = Singular, 3 = 3^rd^ Person, Aux = Auxiliary. Case endings are indicated as follows: NOM = Nominative. The aspect of the verb is marked as follows: PFV = Perfective and IPFV = Imperfective.

^6^ The negativity observed by Wittenberg et al. (2014) pertains to the fact that light verb constructions unlike non-light verb constructions are plausible but have a non-canonical event structure. Coulson and Kutas (2001) observe the negativity in good comprehenders of jokes, which indexes frame shifting. Huang et al. (2010) attributed the negativity seen in case of concrete concepts to imagery.

